# BET inhibition induces HEXIM1- and RAD51-dependent conflicts between transcription and replication

**DOI:** 10.1101/374660

**Authors:** Akhil Bowry, Ann Liza Piberger, Eva Petermann

## Abstract

BET bromodomain proteins are epigenetic readers required for oncogenic transcription activities, and BET inhibitors have been rapidly advanced into clinical trials. Understanding the effects of BET inhibition on other nuclear processes such as DNA replication will be important for future clinical applications. Here we show that BET inhibition causes replication stress in cancer and non-cancer cells due to a rapid burst in global RNA synthesis and interference of transcription with replication. We identify BRD4 as the main BET inhibitor target in this process and provide evidence that BRD4 inhibition causes transcription-replication interference through release of P-TEFb from its inhibitor HEXIM1, promoting RNA Polymerase II phosphorylation. Unusually, BET inhibitor-induced transcription-replication interference does not activate the classic ATM/ATR-dependent DNA damage response. We show however that they promote foci formation of the homologous recombination factor RAD51. Both HEXIM1 and RAD51 are required for BET inhibitor-induced fork slowing, but rescuing fork slowing by HEXIM1 or RAD51 depletion activate a DNA damage response. Our data support a new mechanism where BRD4 inhibition slows replication and suppresses DNA damage through concerted action of transcription and homologous recombination machineries. They shed new light on the roles of DNA replication and recombination in the action of this new class of cancer drugs.

## INTRODUCTION

The BET family of bromodomain containing proteins are epigenetic readers that bind to lysine-acetylated histone tails and regulate transcription by recruiting and activating Positive Transcription Elongation Factor b (P-TEFb). P-TEFb can occur in an inactive complex with 7SK-snRP, and two active complexes with BET protein BRD4 or the super elongation complex (SEC). BRD4 activates P-TEFb by releasing it from 7SK-snRP, and recruits active P-TEFb to gene promoters (Quaresma et al., 2016). Active P-TEFb facilitates RNA Polymerase II (Pol II) pause release by phosphorylating the RNA Pol II C-terminal domain (CTD) and other targets.

BET proteins are required for oncogenic transcriptional programmes, and specific small-molecule inhibitors of BET bromodomains promise a targeted cancer treatment (Delmore et al., 2011). BET inhibition down-regulates MYC protein levels and kills tumour cells in a p53-independent manner (Da Costa et al., 2013). In solid tumour cells however, BET inhibitor responses can be *MYC* independent (Lockwood et al., 2012). Even though the molecular mechanisms surrounding BET inhibitor action are still poorly understood, BET inhibitors are already undergoing clinical trials in a wide range of cancers (Andrieu et al., 2016; Fujisawa and Filippakopoulos, 2017).

More recently, BRD2 and BRD4 have been implicated in DNA replication and DNA damage responses (Da Costa et al., 2013; Floyd et al., 2013; Sansam et al., 2018). BRD4 in particular interacts with DNA replication factors RFC, TICRR and CDC6 (Maruyama et al., 2002; Sansam et al., 2018; Zhang et al., 2018). Inhibiting the interaction between BRD2/4 and TICRR slowed euchromatin replication, and it was suggested that BET proteins control DNA replication initiation to prevent interference between replication and transcription (Sansam et al., 2018). BET inhibitors cause little or no DNA damage, but promote down-regulation of DNA replication stress response and-repair genes (Pawar et al., 2018; Zhang et al., 2018). It is not known whether the latter are specific responses to BET inhibition affecting replication and repair. Investigating more direct effects of BET proteins and BET inhibition on DNA replication might help understand BET inhibitor action independently of cell type-specific transcription programmes and provide new insights into potential side effects and resistance mechanisms.

We previously reported that JQ1 treatment causes replication fork slowing in NALM-6 leukaemia cells, indicative of replication stress (Da Costa et al., 2013). Replication stress occurs when the transcription machinery or other obstacles hinder replication fork progression, which promotes formation of mutagenic or cytotoxic DNA damage, especially double-strand breaks (DSBs). This is highly relevant to cancer therapy, as many conventional chemotherapies act by causing severe replication stress and collapse of replication forks into DSBs. However, non-toxic levels of replication stress can promote genomic instability, an unwanted side effect of cancer therapy (Kotsantis et al., 2015).

Here we describe a new mechanism by which BET inhibition causes replication stress. We show that BET inhibition, and loss of BRD4, cause rapid up-regulation of RNA synthesis and transcription-dependent replication fork slowing in a pathway that depends on HEXIM1 and RAD51. Unexpectedly, combination of BET inhibitor with HEXIM1 or RAD51 depletion prevents fork slowing but activates a DNA damage response, suggesting that replication fork slowing might help suppress BET inhibitor-induced DNA damage.

## RESULTS

U2OS osteosarcoma cells were used as a well-characterised model for replication stress and DNA damage. Osteosarcoma is one of many cancers that have been proposed to benefit from BET inhibitor treatment (Lamoureux et al., 2014). We confirmed that JQ1 treatment slowed replication as early as 1 h after drug addition (Fig. 1A, B). Replication was also slowed by lower concentrations of JQ1 and another BET inhibitor, I-BET151 (Fig. S1A, B).

**Figure 1.**
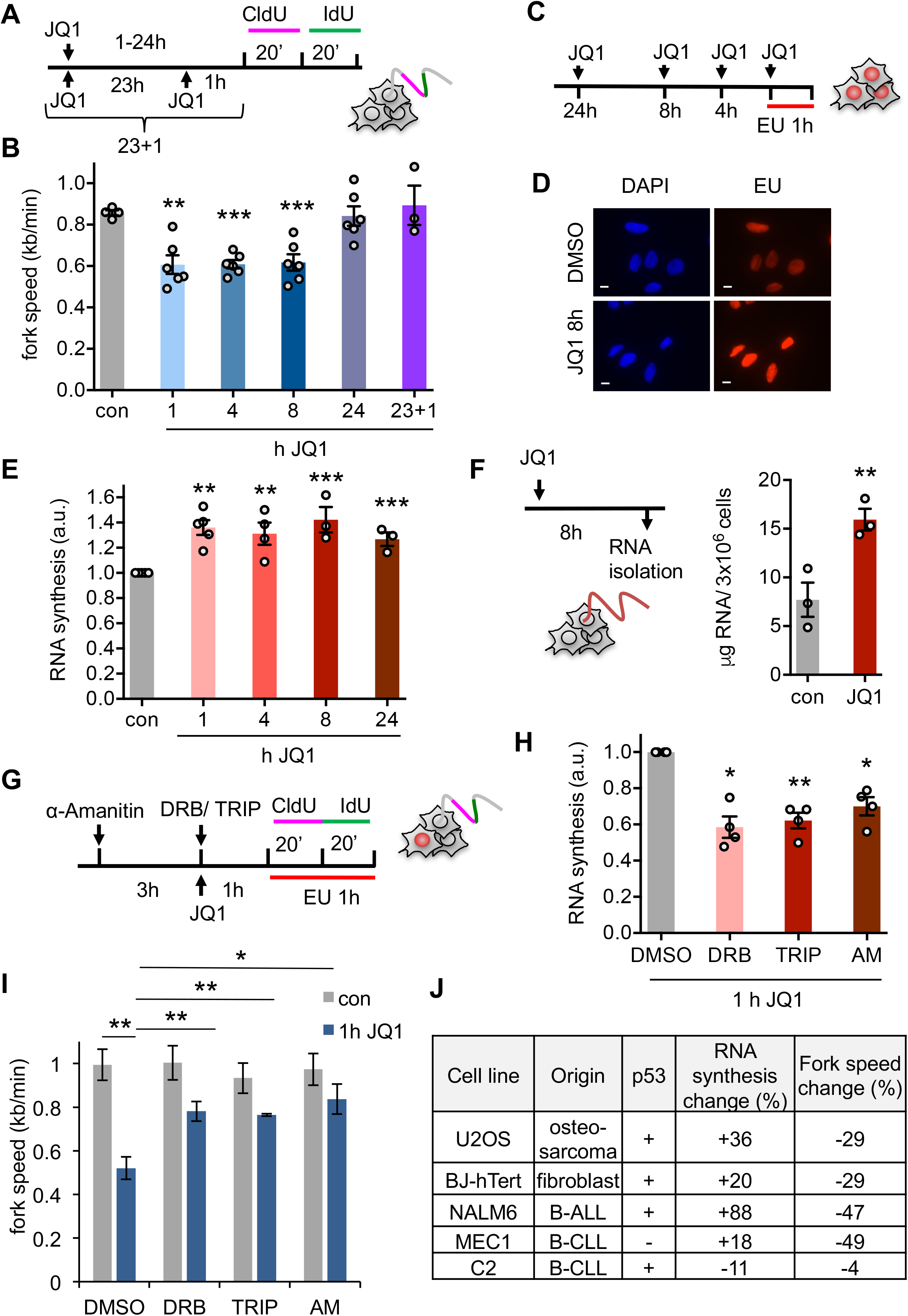
BET inhibition induces replication-transcription conflicts. A) DNA fibre labelling was performed in U2OS cells treated with 1 μM JQ1 for 1-24 h. B) Replication fork speeds after JQ1 treatment. C) U2OS cells were EU-labelled for 1 h after JQ1 treatment. D) Representative images of click-stained EU labelled cells +/- 8 h JQ1. E) Quantification of nuclear EU intensity after JQ1 treatment. F) RNA was extracted after treatment with JQ1 for 8 h, and yield normalised to cell number. G) Cells were treated with transcription inhibitors before and during EU or DNA fibre labelling. H) EU quantification of nascent RNA synthesis in cells treated with transcription inhibitors and JQ1. I) Replication fork speeds after 1h JQ1 treatment +/-transcription inhibitors. J) JQ1 effect on nascent RNA synthesis and replication fork speeds in a panel of human cell lines. Bars: 10 μm.

As reported previously in NALM-6 cells (Da Costa et al., 2013), replication forks speeds were recovered to control levels after 24 h incubation with JQ1. We confirmed that this was not due to loss of JQ1 activity, since re-adding fresh JQ1 after 23 h did not slow fork speeds (Fig. 1A, B). This suggests that replication forks are rapidly slowed by JQ1 treatment, but eventually adapt. Cell cycle distribution remained unaffected between 1-8 h JQ1 treatment, but cells started to accumulate in G1 after 24 h JQ1 treatment (Fig. S1C). The lack of S phase arrest could be explained by compensatory new origin firing (Fig. S1D).

To investigate whether ongoing transcription contributes to JQ1-induced replication slowing, we quantified nascent RNA synthesis using nuclear incorporation of 5-ethynyluridine (EU) (Fig. 1C). EU incorporation increased by around 35% after 1 h JQ1 treatment and throughout the time course (Fig. 1D, E). Increased RNA synthesis was also observed in NALM-6 cells and in U2OS cells treated with I-BET151 (Fig. S2A, B). For an alternative approach, we isolated total RNA and normalised yields to cell numbers, showing that JQ1-treated cells contained more RNA overall (Fig. 1F).

To test whether replication fork slowing was transcription-dependent, we transiently inhibited RNA synthesis using transcription inhibitors triptolide, DRB and α-amanitin (Fig. 1G). These treatments inhibit on-going RNA synthesis and increased replication fork speeds specifically in presence of JQ1 (Fig. 1H, I). Similar results were observed in BJ–hTert fibroblasts, an immortalised human non-cancer cell line (Fig. S2C, D). These data suggest that JQ1-induced replication stress depends on active RNA synthesis and this is not restricted to cancer cells.

We further tested the relationship between increased RNA synthesis and replication fork slowing in two additional chronic leukaemia cell lines, C2 and MEC1 (Fig. S2E-H). In summary, 1 h JQ1 treatment increased RNA synthesis in four out of five cell lines tested, and this was always accompanied by fork slowing (Fig. 1J). Fork slowing was more dramatic in leukaemia lines compared to U2OS or fibroblasts. Only C2 cells appeared resistant to JQ1 effects, displaying neither increased RNA synthesis nor fork slowing.

We used siRNA to investigate which BET protein was the target of JQ1 induced replication-transcription conflicts. We first depleted BRD4, which interacts with DNA replication proteins RFC, TICRR and CDC6 (Maruyama et al., 2002; Sansam et al., 2018; Zhang et al., 2018). BRD4 depletion increased RNA synthesis (Fig. 2A-D) and reduced fork speeds (Fig. 2E). Adding JQ1 did not further affect fork speeds in BRD4-depleted cells (Fig. 2E). Ectopic expression of the long isoform of BRD4 (Fig. 2F, G) and short transcription inhibitor treatments rescued BRD4 siRNA-induced fork slowing (Fig. 2H). In contrast, depletion of BRD2 or BRD3 did not increase RNA synthesis and caused negligible fork slowing (Fig. 2I-N). These data suggest that BRD4 is the BET protein that prevents replication-transcription conflicts and is required for BET inhibitor-induced replication stress.

**Figure 2.**
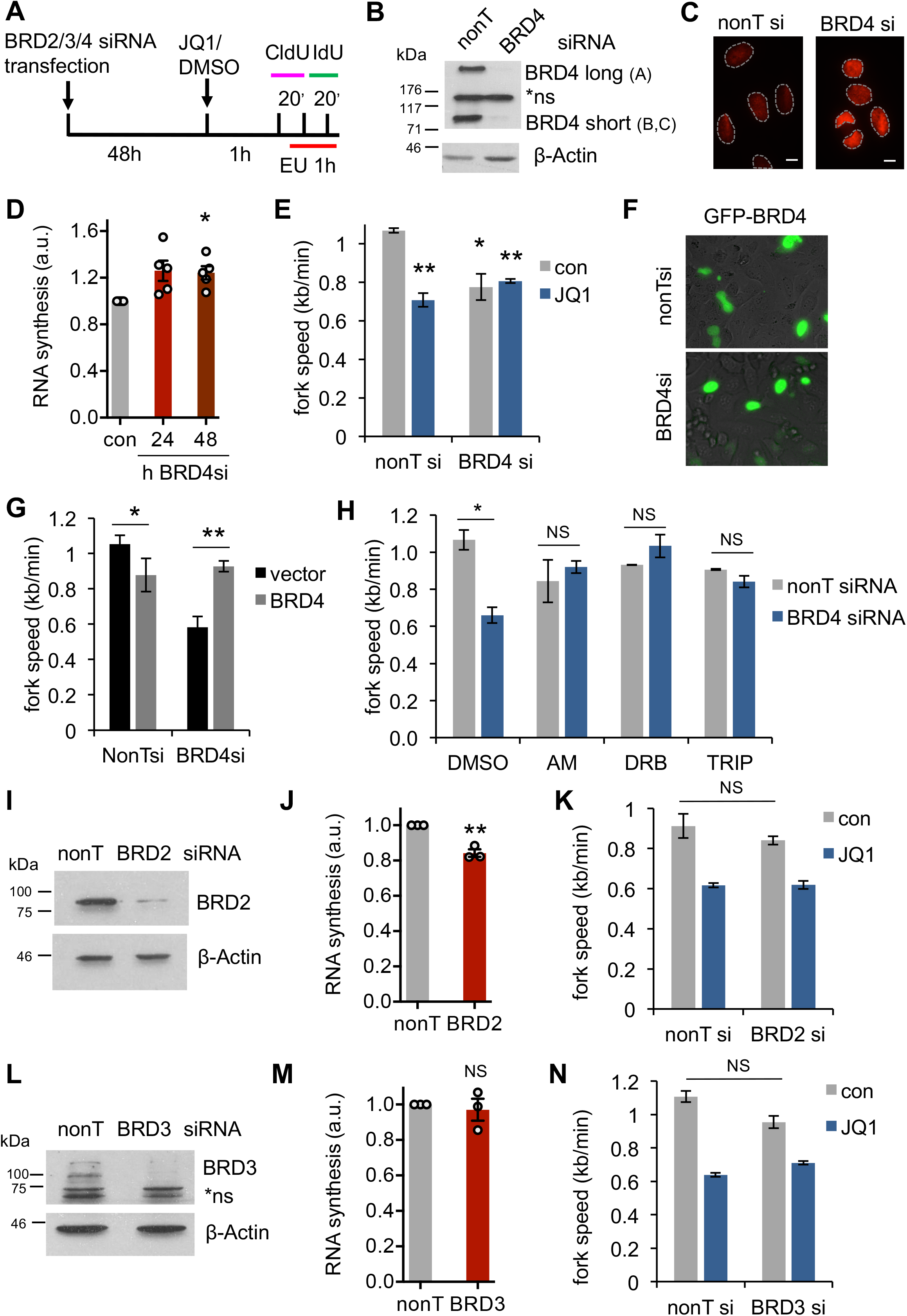
Loss of BRD4 causes replication-transcription conflicts. A) DNA fibre labelling was performed after 48 h BRD2/3/4 depletion. B) Protein levels of BRD4 isoforms after siRNA depletion. NS = non-specific bands. C) Representative images of EU staining after BRD4 siRNA. D) Quantification of nuclear EU intensity after BRD4 siRNA. E) Replication fork speeds after BRD4 depletion +/- 1h JQ1. F) EmGFP-BRD4 expression 48 h after plasmid transfection is similar +/- BRD4 siRNA. G) Replication fork speeds after BRD4 siRNA +/- BRD4 long isoform expression plasmid. H) Median replication fork speeds after BRD4 siRNA +/- transcription inhibitors. I) Protein levels of BRD2 after siRNA depletion. J) Quantification of nuclear EU intensity after BRD2 siRNA. K) Replication fork speeds after BRD2 siRNA +/- 1h JQ1. L) Protein levels of BRD3 after siRNA depletion. NS = non-specific bands. M) Quantification of nuclear EU intensity after BRD3 siRNA. N) Replication fork speeds after BRD3 siRNA +/- 1 h JQ1. Bars: 10 μm.

We then investigated the mechanism of increased RNA synthesis. It has been reported that both JQ1 and I-BET151 can change the equilibrium between the different P-TEFb complexes, disrupting the 7SK-snRP and increasing the proportion of active P-TEFb in complex with the SEC (Bartholomeeusen et al., 2012; Chaidos et al., 2014). This increases transcription of a number of genes such as HEXIM1 (Bartholomeeusen et al., 2012; Rathert et al., 2015). Increased HEXIM1 levels eventually re-establish P-TEFb inhibition (Fig. 3A). We hypothesised that JQ1-induced replication-transcription conflicts could result from 7SK-snRP dissociation and increased RNA Pol II activity. In line with this, RNA Pol II CTD Serine2 phosphorylation was transiently increased during the first hours of JQ1 treatment (Fig. 3B, C). In addition to BET inhibitors, other drugs including hexamethylene bis-acetamide (HMBA) have been reported to dissociate 7SK-snRP from P-TEFb (Fujinaga et al., 2015). If 7SK-snRP dissociation underlies BET inhibitor-induced fork slowing, then HMBA treatment should also slow replication, even though HMBA has no known connection to replication forks. Indeed, HMBA treatment slightly increased RNA synthesis and caused replication fork slowing, which was not further exacerbated by co-treatment with JQ1 (Fig. 3D, E). Finally, the short isoform of BRD4, which cannot effectively interact with P-TEFb (Schroder et al., 2012), failed to rescue the effect of BRD4 knockdown on replication fork slowing (Fig. 3F-H).

**Figure 3.**
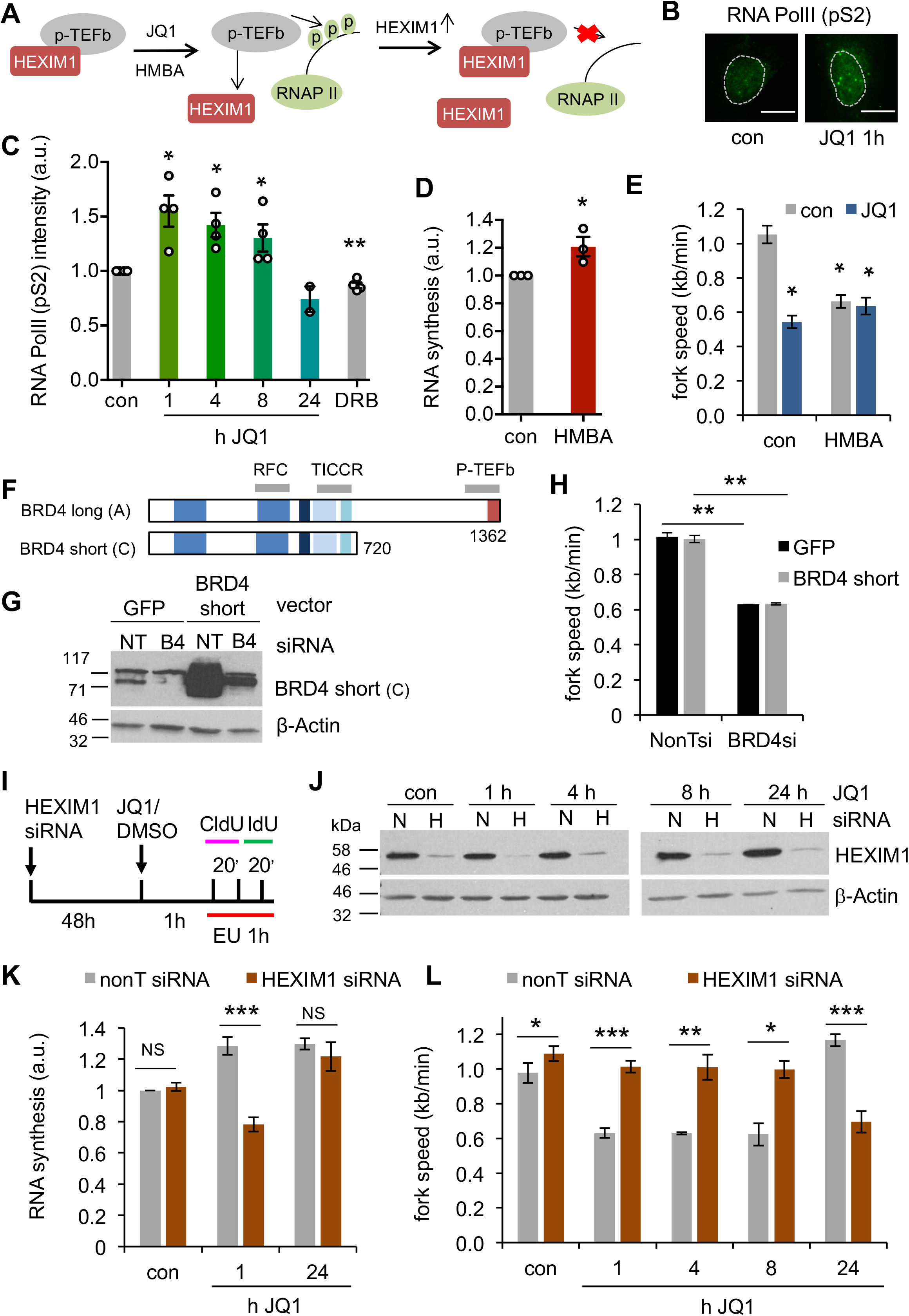
BET inhibitor-induced replication-transcription conflicts require HEXIM1. A) Current model of the role of HEXIM1 in JQ1-induced transcription increase. B) Nuclear immunostaining for phospho-S2 RNA Pol II after JQ1 treatment. C) Quantification of nuclear phospho-S2 RNA Pol II staining as in B. D) Quantification of nuclear EU intensity +/- 5 mM HMBA (1h). E) Replication fork speeds after HMBA treatment +/- JQ1 (1h). F) Schematic of BRD4 isoforms used. G) Protein levels of BRD4 short isoform (C) after full length BRD4 siRNA +/- BRD4 short isoform expression plasmid. H) Replication fork speeds after full length BRD4 siRNA +/- BRD4 short isoform expression plasmid. I) EU and DNA fibre labelling was performed after HEXIM1 depletion for 48 h. J) Protein levels of HEXIM1 after siRNA transfection. K) Quantification of nuclear EU intensity after HEXIM1 siRNA and JQ1 treatment. L) Replication fork speeds after HEXIM1 siRNA and JQ1 treatment.

We decided to further investigate the roles of HEXIM1 in JQ1-induced RNA synthesis and replication fork slowing. HEXIM1 depletion delayed the effects of JQ1, preventing the early increase in RNA synthesis and replication fork slowing. After 24 h JQ1 treatment however, RNA synthesis increased, accompanied by fork slowing (Fig. 3I-L). These data suggest that early JQ1-induced RNA synthesis and transcription-replication conflicts and replication recovery after 24 h JQ1 both require HEXIM1. However, we observed no increase in HEXIM1 protein levels or decrease in nascent RNA synthesis levels during 24 h JQ1 treatment, suggesting that the process of adaptation is more complex than a HEXIM1-mediated feedback loop suppressing transcription (Fig. 3J).

We then investigated the relationship between JQ1-induced fork slowing and DNA damage response. Replication fork slowing can expose single-stranded DNA (ssDNA) and, if forks collapse, cause DSBs. These activate the ATR and ATM checkpoint kinases and p53. However, JQ1 does not activate p53 (Da Costa et al., 2013) or induce DNA damage (Pawar et al., 2018). In line with this, we observed no JQ1-induced increase in ssDNA or DSBs as measured by nuclear foci formation of RPA and 53BP1, or phosphorylation of the ATM and ATR targets histone H2AX (γH2AX, Fig. 4A), CHK1 or RPA32 (Fig. S3A). BRD4 depletion also failed to induce DNA damage foci (Fig. S3B).

**Figure 4.**
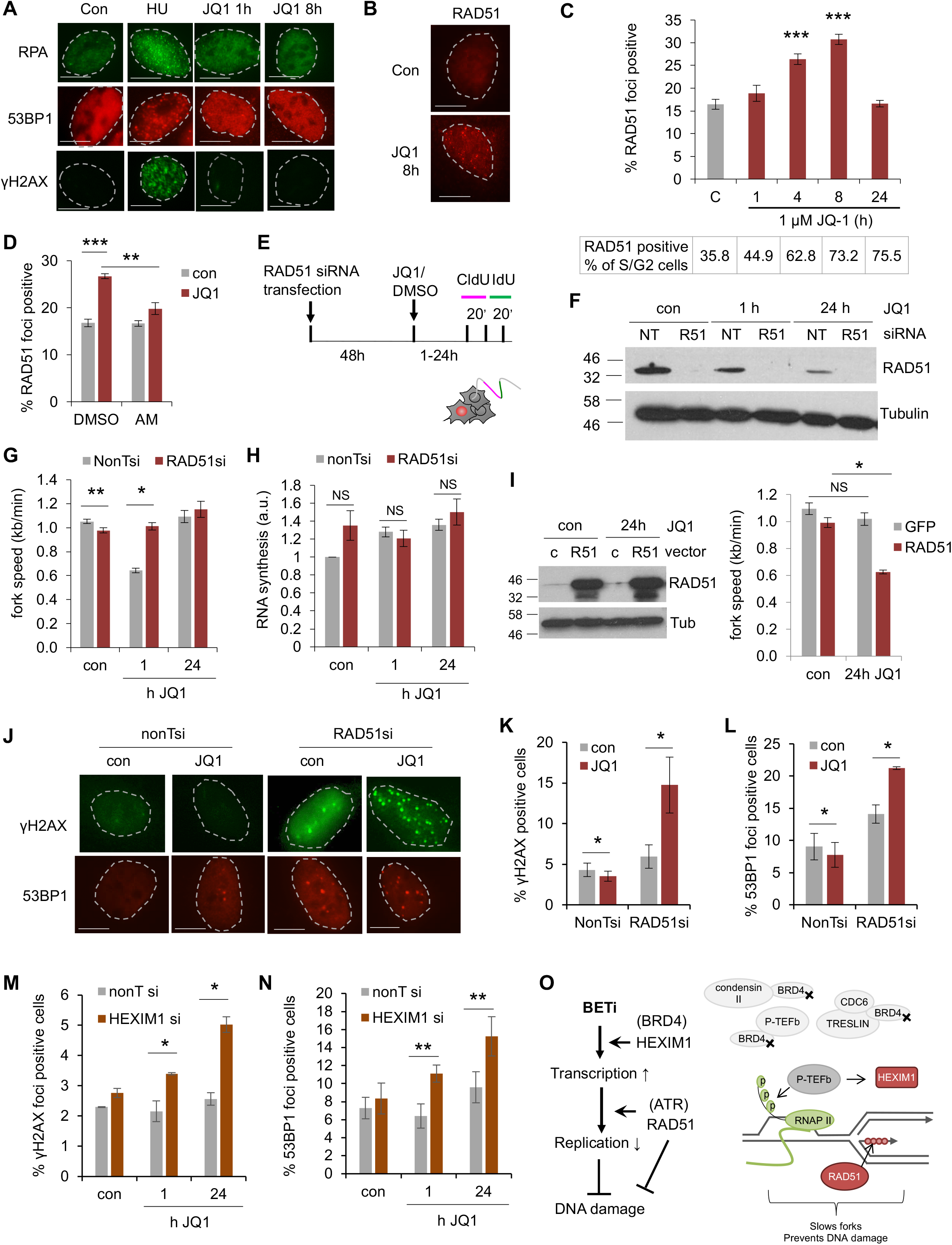
RAD51 and HEXIM1 modulate BET inhibitor-induced DNA damage. A) Representative images of RPA, γH2AX and 53BP1 foci after JQ1 treatment. 2 mM HU, 8 h, was used as positive control. B) Representative images of RAD51 foci after JQ1 treatment. C) Total percentages (top) and percent normalised to S/G2 content (bottom) of cells containing more than 5 RAD51 foci after JQ1 treatment. D) Percentages of cells containing more than 5 RAD51 foci +/- 4 h JQ1 and *α*-amanitin (AM). E) EU and DNA fibre labelling was performed after 48 h RAD51 depletion. F) Protein levels of RAD51 after siRNA depletion. G) Replication fork speeds after RAD51 siRNA and JQ1 treatment. H) Quantification of nuclear EU intensity +/- RAD51 siRNA and JQ1 treatment. I) Levels of RAD51 and loading control, and replication fork speeds after transient overexpression of RAD51 or eGFP (control) for 24 h +/- JQ1. J) Representative images of γH2AX and 53BP1 foci after JQ1 treatment +/- RAD51 siRNA. K) Percentages of cells containing more than 8 γH2AX foci after JQ1 +/- RAD51 siRNA. L) Percentages of cells containing more than 8 53BP1 foci after JQ1 +/- RAD51 siRNA. M) Percentages of cells containing more than 8 γH2AX foci after JQ1 +/- HEXIM1 siRNA. N) Percentages of cells containing more than 8 53BP1 foci after JQ1 +/- HEXIM1 siRNA. O) Model: BET inhibition disrupts recruitment of chromatin and replication factors; increased RNA synthesis and RAD51 activity (e.g. fork reversal) slow replication forks without generating DNA damage. Bars: 10 μm.

Unexpectedly however, JQ1 treatment induced foci formation of the homologous recombination factor RAD51, which depended on ongoing transcription and ATR activity (Fig. 4B-D, S3C). This suggested that RAD51 is recruited in response to JQ1-induced transcription-replication conflicts, aided by a basal level of ATR activity. We used siRNA to investigate the impact of RAD51 on replication fork progression in JQ1-treated cells (Fig. 4E). Interestingly, RAD51 depletion prevented JQ1-induced fork slowing (Fig. 4F, G). This phenomenon, previously reported for DNA damaging treatments, is suggestive of RAD51 recruitment to forks and fork reversal (Zellweger et al., 2015). We investigated whether RAD51 suppresses RNA synthesis, like HEXIM1. RAD51-depleted cells displayed high levels of RNA synthesis both in presence and absence of JQ1, making these data difficult to interpret (Fig. 4H). Nevertheless, we concluded that RAD51-dependent rescue of fork speeds was not due to global loss of transcription. Transient overexpression of RAD51 was able to promote fork slowing after 24 h JQ1 treatment, additionally supporting that RAD51 directly slows forks in response to JQ1 (Fig. 4I). We then tested the effect of RAD51 depletion on JQ1-induced DNA damage. Interestingly, despite rescuing fork progression, RAD51 depletion actually promoted DNA damage as indicated by foci formation of γH2AX and 53BP1 (Fig. 4J-L). This suggests that RAD51 is a major factor preventing DNA damage at JQ1-slowed replication forks.

Because HEXIM1 depletion had also rescued fork progression, we tested the effect of HEXIM1 depletion on DNA damage. Indeed, HEXIM1-depleted cells accumulated DNA damage early during JQ1 treatment that persisted after 24 h JQ1 (Fig. 4M, N). In agreement with the increased DNA damage, HEXIM1-depleted cells were more sensitive to 24 h JQ1 treatment than control cells, as were ATR inhibitor-treated cells (Fig. S3D). This suggests that both HEXIM1, which is upstream of transcription-replication conflicts, and RAD51, which acts downstream of these conflicts, slow replication forks and prevent JQ1 from inducing DNA damage (Fig. 4O).

## DISCUSSION

We report that BET inhibitor-induced P-TEFb activation causes transcription-dependent replication fork slowing. This process depends on HEXIM1, which is a central factor in the BET inhibitor response, and the homologous recombination factor RAD51, which is central in the replication stress response. Unusually, BET inhibitor-induced replication stress is transient and, despite engaging RAD51, does not activate a full DNA damage response. We speculate that these unusual transcription-replication conflicts will help illuminate some aspects of BET inhibitor treatment responses. Intriguingly, loss of both HEXIM1 and RAD51 have been implicated in BET inhibitor resistance.

Our data provide new insight into the roles of BRD4 in DNA replication. It has been reported that BRD4 interacts with DNA replication factors RFC, TICRR and CDC6 (Maruyama et al., 2002; Sansam et al., 2018; Zhang et al., 2018). We show here that BRD4 also regulates DNA replication via its P-TEFb interaction. The BRD4 short isoform failed to rescue replication stress, despite containing the interaction domains for TICRR and RFC (Maruyama et al., 2002; Sansam et al., 2018). The interaction with CDC6 has not been mapped (Zhang et al., 2018). Our data do not suggest that TICRR and RFC are involved in the phenotypes described here. Instead they support previous reports that short-term BET inhibitor treatments are associated with increased P-TEFb activity (Bartholomeeusen et al., 2012; Chaidos et al., 2014), and we show that this also involves increased RNA synthesis. We previously reported that overexpression of oncogene H-RAS^V12^, or transcription factor TBP increase nascent RNA synthesis and cause replication stress (Kotsantis et al., 2016). BET inhibitors present a way in which small molecule inhibitor treatments can cause replication stress by increasing RNA synthesis. Any treatments that disrupt the complex of P-TEFb with 7SK-snRP, including other cancer drugs such as HDAC inhibitors and azacytidine (Fujinaga et al., 2015), could potentially cause transcription-replication conflicts through this pathway.

Strikingly, our findings suggest that HEXIM1 is required for BET inhibitor-induced replication fork slowing, which is delayed by at least 8 hours in absence of HEXIM1. After 24 hours, JQ1-induced gene expression changes or possible compensation with HEXIM2 (Byers et al., 2005) will make these later effects harder to interpret. HEXIM1 depletion has been associated with long-term BET inhibitor resistance. Our findings do not conflict with these reports, as BET inhibitor resistance in HEXIM1-depleted cells was only observed after prolonged (>24 hour) treatment (Devaraj et al., 2016). It was proposed that HEXIM1 loss decreases overall JQ1 effectiveness by allowing higher P-TEFb activity (Devaraj et al., 2016). Importantly, we show that HEXIM1 depletion can actually promote JQ1 effects, such as DNA damage. In addition to modulating transcription-dependent fork slowing, HEXIM1 might play undiscovered roles in replication stress and DNA damage response. It may be relevant that HEXIM1 also regulates p53 (Lew et al., 2012). Nevertheless, we observed JQ1-induced replication fork slowing in a p53 mutant cell line. Highly variable HEXIM1 protein levels have been observed in cancers, including reduced HEXIM1 levels in metastatic breast cancer (Ketchart et al., 2013), melanoma (Tan et al., 2016), and acute leukaemia (Devaraj et al., 2016; Huang et al., 2016). It will be important for cancer researchers to further investigate the relationship between HEXIM1, replication stress, and the response to cancer treatment.

Homologous recombination capacity in cancer cells is a well-established predictive biomarker for the response to replication stress-inducing treatments. RAD51 loading is known to actively slows forks in response to a variety of genotoxic agents (Zellweger et al., 2015). We show that BET inhibition also activates RAD51, likely due to increased transcription, and that RAD51 promotes fork slowing in response to BET inhibitor. RAD51 may slow forks at sites of direct collisions between replication and transcription machineries, or perhaps also in response to indirect effects of transcription on replication. RAD51 expression is down-regulated in response to BET inhibition (Yang et al., 2017) and in models of acquired BET inhibitor resistance (Pawar et al., 2018). Intriguingly, acquired BET inhibitor resistance models also displayed increased DNA damage signalling (Pawar et al., 2018). This agrees with our data that RAD51 downregulation increases DNA damage signalling.

Our data support a speculative model where loss of HEXIM1 or RAD51 allows normal replication fork progression in the presence of BET inhibitor, but at the expense of DNA damage. Such a process may even contribute to long–term BET inhibitor resistance. While it is extensively documented that transcription-replication conflicts promote DNA damage, BET inhibition seems to induce DNA damage when such conflicts are prevented. This damage might e.g. result from reduced chromatin recruitment of chromatin remodellers (Floyd et al., 2013) or DNA replication proteins (Sansam et al., 2018; Zhang et al., 2018) (Fig. 4O). The underlying mechanisms, and how they relate to BET inhibitor response, will require future investigation. In summary, we provide new insights into the relationship between BET proteins, DNA replication and DNA damage response that will be relevant to cancer research.

## EXPERIMENTAL PROCEDURES

### DNA fibre analysis

Cells were labelled with 25 μM CldU and 250 μM IdU for 20 min each and DNA fibre spreads prepared. HCl-treated fibre spreads were incubated with rat anti-BrdU (BU1/75, Abcam, 1:250) and mouse anti-BrdU (B44, Becton Dickinson, 1:500) for 1 h, fixed with 4% PFA and incubated with anti-rat AlexaFluor 555 and anti-mouse AlexaFluor 488 (Thermo Fisher) for 1.5 h.

### EU incorporation assay

5-ethynyl uridine (EU) incorporation assays were performed using the Click-iT RNA Alexa Fluor 594 Imaging Kit (Invitrogen). Cells were incubated with 1 mM EU for 1 h, followed fixation and Click-iT reaction according to manufacturer’s instructions. DNA was counterstained with DAPI. Nuclear masks were generated in ImageJ to quantify mean fluorescence intensities per nucleus. Results were normalised to control/DMSO to account for variation in staining intensity.

### Statistical analysis

Values are means +/- 1x SEM of at least three independent repeats, unless indicated otherwise. Statistical tests were the one-tailed student’s t-test, or 2- way ANOVA with Tukey’s for colony assays. Unless indicated otherwise, asterisks compare to control and signify *p<0.05, **p<0.01, ***p<0.001.

## Acknowledgements

A.B. and A.L.P were supported by an MRC studentship (1552339) and a German Research Foundation Fellowship (Pl 1300/1-1), respectively. We thank Drs Catherine Rogers and Panagis Filippakopoulos for BRD4 expression constructs.

## Author contributions

A.B. designed and performed experiments, analysed data and contributed to writing the paper; A.L.P performed RAD51 foci quantification, E.P. conceived the project, designed experiments and wrote the paper.

## Supplemental Figures

**Figure S1.**
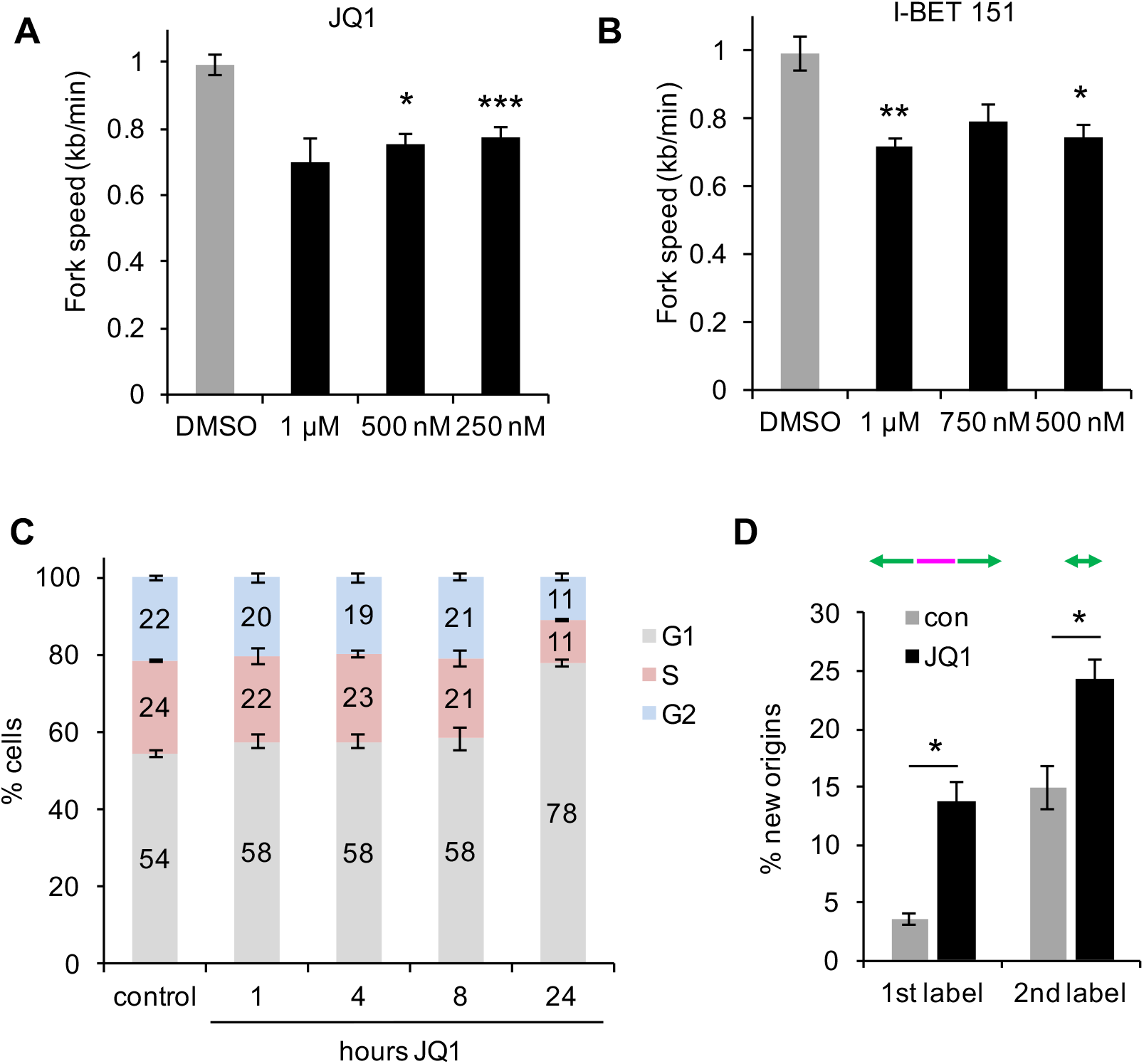
BET inhibition induces replication fork slowing. A) Replication fork speeds in U2OS cells after treatment with 0.25-1 μM JQ1 for 1 h. B) Replication fork speeds in U2OS cells after treatment with 0.5-1 μM I-BET151 for 1 h. C) Cell cycle distribution measured by flow cytometry of U2OS cells after 1-24 h treatment with 1 μM JQ1. D) New origin firing in U2OS cells after treatment with 1 μM JQ1 for 1 h.

**Figure S2.**
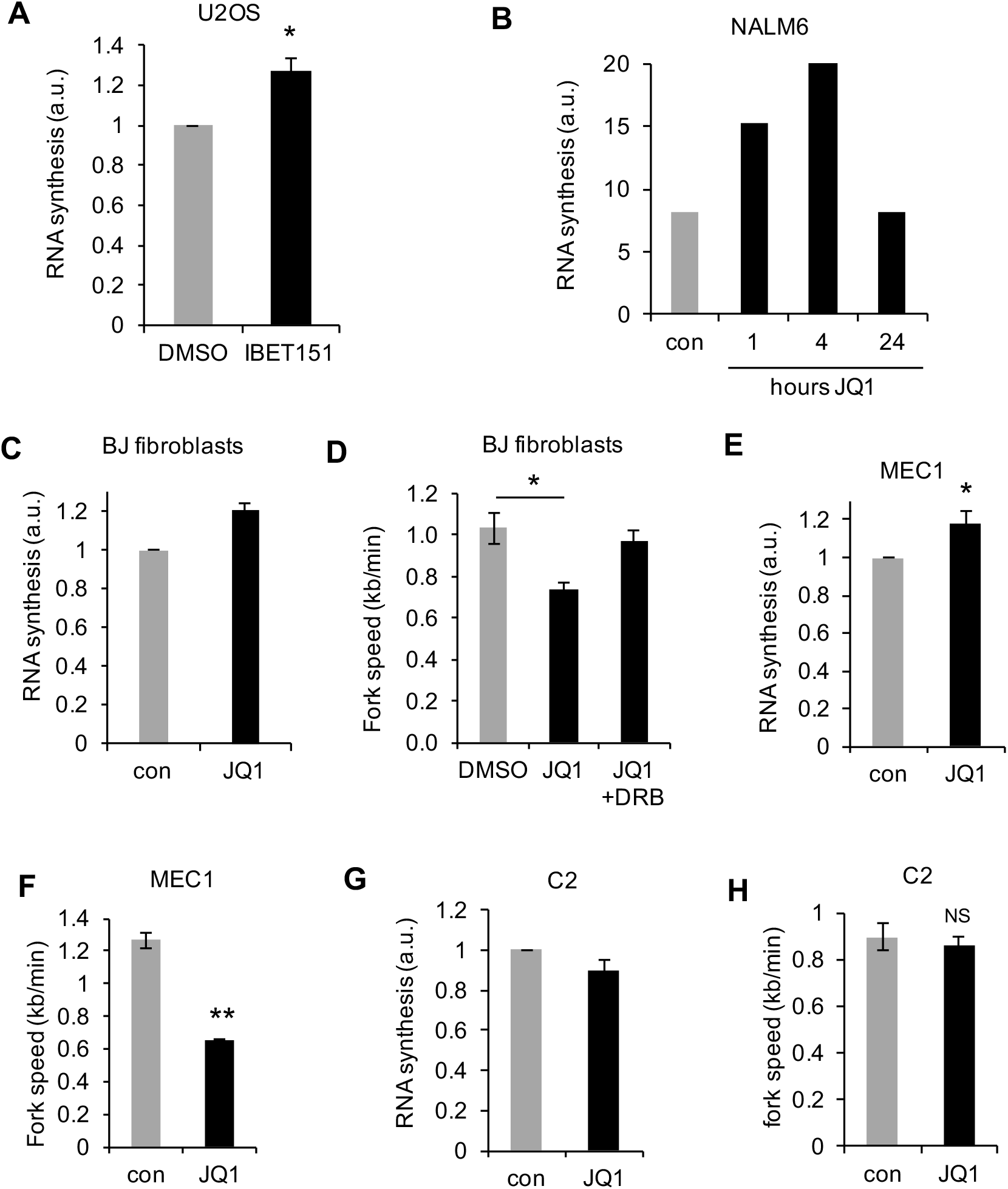
Replication-transcription conflicts induced by I-BET151 and in a range of cell lines. A) Quantification of nuclear EU intensity in U2OS cells after I-BET151 (1h). B) Nuclear EU intensity in NALM6 cells after 1-24h 1 μM JQ1. C) Nuclear EU intensity in BJ-hTert fibroblasts +/- 1 μM JQ1 (1h). D) Median replication fork speeds in BJ-hTert fibroblasts +/- 1 μM JQ1 (1h), and treated with JQ1 and DRB for 1h. E) Nuclear EU intensity in MEC1 cells +/-1 μM JQ1 (1h). F) Median replication fork speeds in MEC1 cells +/- 1 μM JQ1 (1h). G) Nuclear EU intensity in C2 cells +/-1 μM JQ1 (1h). H) Median replication fork speeds in C2 cells +/- 1 μM JQ1 (1h).

**Figure S3.**
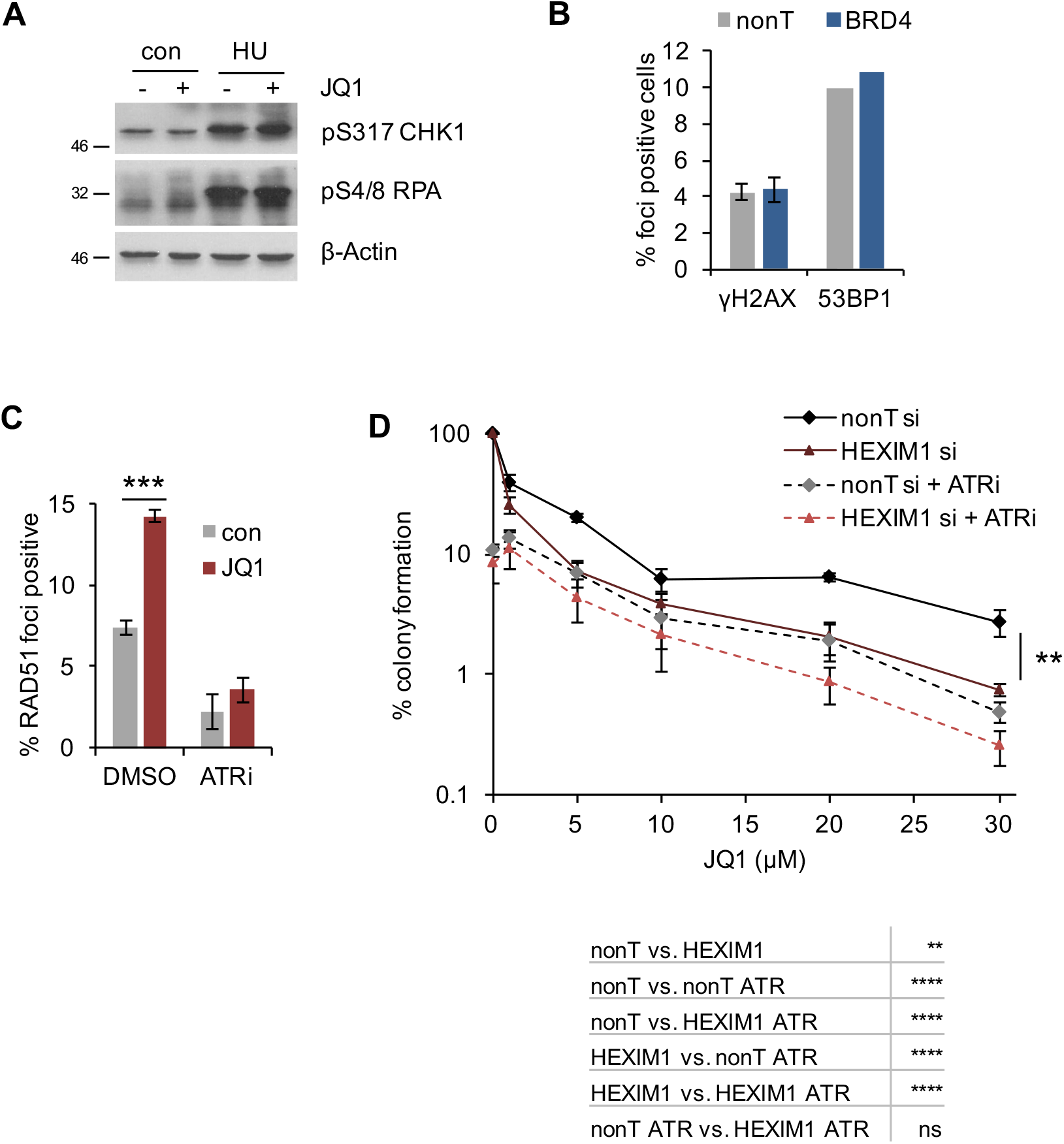
DNA damage response and cell death after BET inhibition. A) Levels of phospho-S4/8 RPA32, phospho-S317 CHK1 and β-Actin after treatment with JQ1 and HU as indicated. B) Percentages of cells containing more than 8 γH2AX or 53BP1 foci after BRD4 depletion. N = 2 (γH2AX), N = 1 (53BP1). C) Percentages of cells containing more than 8 RAD51 foci after JQ1 treatment +/- ATR inhibitor AZ20. D) Colony survival of U2OS cells treated with indicated concentrations of JQ1 for 24h, +/- HEXIM1 siRNA and AZ20. The statistical test used was 2-way ANOVA with Tukey’s, **** = p<0.0001.

## Supplemental Experimental Procedures

### Cell lines and reagents

Human U2OS, NALM-6, C2, MEC1 and BJ-hTert cells (ATCC) were authenticated using 8-locus STR profiling (LGC Standards). Cells were confirmed Mycoplasma free and grown in DMEM (high glucose) or RPMI 1640 in a humidified atmosphere containing 5% CO_2_.

JQ1 (1 uM), I-BET151 (Dawson et al., 2011), AZ20 (2.4 μM) and triptolide (1 μM) were from Tocris Bioscience. 5,6-dichloro-1-β-D-ribofuranosyl-1H-benzimidazole (DRB, 100 μM), α-amanitin (10μg/ml), hydroxyurea (2 mM), and camptothecin were from Sigma.

### siRNA and DNA transfection

siRNAs against BRD2, BRD3, BRD4, HEXIM1 (L-004935-00, L-004936-00, L-004937-00, L-012225-01-0005) and RAD51 (Ito et al., 2005), were from Dharmacon, and “Allstars negative control siRNA” from Qiagen. Cells were transfected with 50 nM siRNA using Dharmafect 1 reagent (Dharmacon). For BRD4 and RAD51 overexpression cells were transfected using TransIT-2020 (Mirus Bio) with 2.5 μg pcDNA6.2/N-EmGFP-BRD4(long)-DEST (Philpott et al., 2014), pCDNA5-3HA-BRD4(short)-DEST (kind gift from Dr Panagis Filippakopoulos), pcDNA3.1/V5/His-TOPO-RAD51 (Sorensen et al., 2005), or control plasmids pcDNA3.1(+) (Invitrogen) or pEGFP-C2 (Clontech).

### Immunofluorescence

Cells were fixed with 4% PFA for 10 min and blocked with 2% BSA. Primary antibodies were mouse anti-phospho-Histone H2AX (Ser139) (JBW301, Millipore 05-636, 1:1,000), rabbit anti-53BP1 (Bethyl A300-272A, 1:30,000), anti-RNA Pol II pS2 (abcam, 1:500, ab24758), anti-RPA32 (Merck, NA18,1:500) and anti-RAD51 (Abcam, ab63801, 1:500). Secondary antibodies were anti-mouse IgG AlexaFluor 488 and anti-rabbit IgG AlexaFluor 555 (Molecular Probes). DNA was counterstained with DAPI and images were acquired on a Nikon E600 microscope with a Nikon Plan Apo 60x (1.3 NA) oil lens, a Hamamatsu digital camera (C4742-95) and the Volocity acquisition software (Perkin Elmer). Images were analysed using ImageJ (Schneider et al., 2012).

### Western blotting

Cell extracts were prepared in UTB buffer (50 mM Tris-HCl pH 7.5, 150 mM β-mercaptoethanol, 8 M urea) and sonicated to release DNA-bound proteins. Primary antibodies used were rabbit anti-BRD2 (Abcam ab139690, 1:1000), rabbit anti-BRD3 (Abcam ab83478,1:500), rabbit anti-BRD4 (Abcam, ab128874, 1:1000), rabbit anti-HEXIM1 (Abcam ab25388,1:5000), rabbit anti-pS4/S8 RPA32 (Bethyl, A300-245A,1:1000), rabbit anti-pS317 CHK1 (New England Biolabs, 2334S 1:1,000), anti-RAD51 (Abcam, ab63801, 1:1000) mouse anti-PARP1 (F-22, Santa Cruz sc8007, 1:1000), mouse anti-αTUBULIN (B512, Sigma T6074, 1:10,000), rabbit anti-βACTIN (Cell Signaling A967S, 1:5,000).

### RNA isolation

Cells were harvested and counted, and RNA was isolated using Qiagen RNeasy Mini Kit according to manufacturer’s instructions. RNA concentration was determined using an Implen NanoPhotometer Pearl and normalised to cell numbers.

### Colony survival assay

Defined numbers of cells were plated in duplicate before treatment with JQ1 (1 – 30 μM) for 24 h. Colonies of >50 cells were allowed to form in fresh medium and fixed in 50% ethanol, 2% methylene blue.

### Flow cytometry

Cells were fixed with cold 70% ethanol before staining with 10 μg/ml propidium iodide. Cell cycle profiles were gathered using the BD LSR Fortessa X20 and analysed with BD FacsDiva software.

## References

Andrieu, G., Belkina, A.C., and Denis, G.V. (2016). Clinical trials for BET inhibitors run ahead of the science. Drug Discov Today Technol 19, 45–50.

Bartholomeeusen, K., Xiang, Y., Fujinaga, K., and Peterlin, B.M. (2012). Bromodomain and extra-terminal (BET) bromodomain inhibition activate transcription via transient release of positive transcription elongation factor b (P-TEFb) from 7SK small nuclear ribonucleoprotein. J Biol Chem 287, 36609–36616.

Byers, S.A., Price, J.P., Cooper, J.J., Li, Q., and Price, D.H. (2005). HEXIM2, a HEXIM1-related protein, regulates positive transcription elongation factor b through association with 7SK. J Biol Chem 280, 16360–16367.

Chaidos, A., Caputo, V., Gouvedenou, K., Liu, B., Marigo, I., Chaudhry, M.S., Rotolo, A., Tough, D.F., Smithers, N.N., Bassil, A.K., et al. (2014). Potent antimyeloma activity of the novel bromodomain inhibitors I-BET151 and I-BET762. Blood 123, 697–705.

Da Costa, D., Agathanggelou, A., Perry, T., Weston, V., Petermann, E., Zlatanou, A., Oldreive, C., Wei, W., Stewart, G., Longman, J., et al. (2013). BET inhibition as a single or combined therapeutic approach in primary paediatric B-precursor acute lymphoblastic leukaemia. Blood Cancer J 3, e126.

Delmore, Jake E., Issa, Ghayas C., Lemieux, Madeleine E., Rahl, Peter B., Shi, J., Jacobs, Hannah M., Kastritis, E., Gilpatrick, T., Paranal, Ronald M., Qi, J., et al. (2011). BET Bromodomain Inhibition as a Therapeutic Strategy to Target c-Myc. Cell 146, 904–917.

Devaraj, S.G., Fiskus, W., Shah, B., Qi, J., Sun, B., Iyer, S.P., Sharma, S., Bradner, J.E., and Bhalla, K.N. (2016). HEXIM1 induction is mechanistically involved in mediating anti-AML activity of BET protein bromodomain antagonist. Leukemia 30, 504–508.

Floyd, S.R., Pacold, M.E., Huang, Q., Clarke, S.M., Lam, F.C., Cannell, I.G., Bryson, B.D., Rameseder, J., Lee, M.J., Blake, E.J., et al. (2013). The bromodomain protein Brd4 insulates chromatin from DNA damage signalling. Nature 498, 246–250.

Fujinaga, K., Luo, Z., Schaufele, F., and Peterlin, B.M. (2015). Visualization of positive transcription elongation factor b (P-TEFb) activation in living cells. J Biol Chem 290, 1829–1836.

Fujisawa, T., and Filippakopoulos, P. (2017). Functions of bromodomain-containing proteins and their roles in homeostasis and cancer. Nat Rev Mol Cell Biol 18, 246–262.

Huang, M., Garcia, J.S., Thomas, D., Zhu, L., Nguyen, L.X., Chan, S.M., Majeti, R., Medeiros, B.C., and Mitchell, B.S. (2016). Autophagy mediates proteolysis of NPM1 and HEXIM1 and sensitivity to BET inhibition in AML cells. Oncotarget 7, 74917–74930.

Ketchart, W., Smith, K.M., Krupka, T., Wittmann, B.M., Hu, Y., Rayman, P.A., Doughman, Y.Q., Albert, J.M., Bai, X., Finke, J.H., et al. (2013). Inhibition of metastasis by HEXIM1 through effects on cell invasion and angiogenesis. Oncogene 32, 3829–3839.

Kotsantis, P., Jones, R.M., Higgs, M.R., and Petermann, E. (2015). Cancer therapy and replication stress: forks on the road to perdition. Adv Clin Chem 69, 91–138.

Kotsantis, P., Silva, L.M., Irmscher, S., Jones, R.M., Folkes, L., Gromak, N., and Petermann, E. (2016). Increased global transcription activity as a mechanism of replication stress in cancer. Nature communications 7, 13087.

Lamoureux, F., Baud’huin, M., Rodriguez Calleja, L., Jacques, C., Berreur, M., Redini, F., Lecanda, F., Bradner, J.E., Heymann, D., and Ory, B. (2014). Selective inhibition of BET bromodomain epigenetic signalling interferes with the bone-associated tumour vicious cycle. Nature communications 5, 3511.

Lew, Q.J., Chia, Y.L., Chu, K.L., Lam, Y.T., Gurumurthy, M., Xu, S., Lam, K.P., Cheong, N., and Chao, S.H. (2012). Identification of HEXIM1 as a positive regulator of p53. J Biol Chem 287, 36443–36454.

Lockwood, W.W., Zejnullahu, K., Bradner, J.E., and Varmus, H. (2012). Sensitivity of human lung adenocarcinoma cell lines to targeted inhibition of BET epigenetic signaling proteins. Proc Natl Acad Sci U S A 109, 19408–19413.

Maruyama, T., Farina, A., Dey, A., Cheong, J., Bermudez, V.P., Tamura, T., Sciortino, S., Shuman, J., Hurwitz, J., and Ozato, K. (2002). A Mammalian bromodomain protein, brd4, interacts with replication factor C and inhibits progression to S phase. Mol Cell Biol 22, 6509–6520.

Pawar, A., Gollavilli, P.N., Wang, S., and Asangani, I.A. (2018). Resistance to BET Inhibitor Leads to Alternative Therapeutic Vulnerabilities in Castration-Resistant Prostate Cancer. Cell Rep 22, 2236–2245.

Quaresma, A.J.C., Bugai, A., and Barboric, M. (2016). Cracking the control of RNA polymerase II elongation by 7SK snRNP and P-TEFb. Nucleic Acids Research 44, 7527–7539.

Rathert, P., Roth, M., Neumann, T., Muerdter, F., Roe, J.S., Muhar, M., Deswal, S., Cerny-Reiterer, S., Peter, B., Jude, J., et al. (2015). Transcriptional plasticity promotes primary and acquired resistance to BET inhibition. Nature 525, 543–547.

Sansam, C.G., Pietrzak, K., Majchrzycka, B., Kerlin, M.A., Chen, J., Rankin, S., and Sansam, C.L. (2018). A mechanism for epigenetic control of DNA replication. Genes Dev 32, 224–229.

Schroder, S., Cho, S., Zeng, L., Zhang, Q., Kaehlcke, K., Mak, L., Lau, J., Bisgrove, D., Schnolzer, M., Verdin, E., et al. (2012). Two-pronged binding with bromodomain-containing protein 4 liberates positive transcription elongation factor b from inactive ribonucleoprotein complexes. J Biol Chem 287, 1090–1099.

Tan, J.L., Fogley, R.D., Flynn, R.A., Ablain, J., Yang, S., Saint-Andre, V., Fan, Z.P., Do, B.T., Laga, A.C., Fujinaga, K., et al. (2016). Stress from Nucleotide Depletion Activates the Transcriptional Regulator HEXIM1 to Suppress Melanoma. Mol Cell 62, 34–46.

Yang, L., Zhang, Y., Shan, W., Hu, Z., Yuan, J., Pi, J., Wang, Y., Fan, L., Tang, Z., Li, C., et al. (2017). Repression of BET activity sensitizes homologous recombination-proficient cancers to PARP inhibition. Sci Transl Med 9.

Zellweger, R., Dalcher, D., Mutreja, K., Berti, M., Schmid, J.A., Herrador, R., Vindigni, A., and Lopes, M. (2015). Rad51-mediated replication fork reversal is a global response to genotoxic treatments in human cells. J Cell Biol 208, 563–579.

Zhang, J., Dulak, A.M., Hattersley, M.M., Willis, B.S., Nikkila, J., Wang, A., Lau, A., Reimer, C., Zinda, M., Fawell, S.E., et al. (2018). BRD4 facilitates replication stress-induced DNA damage response. Oncogene.

## Supplemental references

Dawson, M.A., Prinjha, R.K., Dittmann, A., Giotopoulos, G., Bantscheff, M., Chan, W.I., Robson, S.C., Chung, C.W., Hopf, C., Savitski, M.M., et al. (2011). Inhibition of BET recruitment to chromatin as an effective treatment for MLL-fusion leukaemia. Nature 478, 529–533.

Ito, M., Yamamoto, S., Nimura, K., Hiraoka, K., Tamai, K., and Kaneda, Y. (2005). Rad51 siRNA delivered by HVJ envelope vector enhances the anti-cancer effect of cisplatin. J Gene Med 7, 1044–1052.

Philpott, M., Rogers, C.M., Yapp, C., Wells, C., Lambert, J.P., Strain-Damerell, C., Burgess-Brown, N.A., Gingras, A.C., Knapp, S., and Muller, S. (2014). Assessing cellular efficacy of bromodomain inhibitors using fluorescence recovery after photobleaching. Epigenetics & chromatin 7.

Schneider, C.A., Rasband, W.S., and Eliceiri, K.W. (2012). NIH Image to ImageJ: 25 years of image analysis. Nat Methods 9, 671–675.

Sorensen, C.S., Hansen, L.T., Dziegielewski, J., Syljuasen, R.G., Lundin, C., Bartek, J., and Helleday, T. (2005). The cell-cycle checkpoint kinase Chk1 is required for mammalian homologous recombination repair. Nat Cell Biol 7, 195–201.

